# Food and temperature change photoperiodic responses in two vole species: different roles for hypothalamic genes

**DOI:** 10.1101/2021.01.20.427447

**Authors:** Laura van Rosmalen, Roelof A. Hut

## Abstract

Seasonal timing of reproduction in voles is driven by photoperiod. Here we hypothesize that a negative energy balance can modify spring-programmed photoperiodic responses in the hypothalamus, controlling reproductive organ development. We manipulated energy balance by the ‘work-for-food’ protocol, in which voles were exposed to increasing levels of food scarcity at different ambient temperatures under long photoperiod. We reveal that common (*Microtus arvalis*) and tundra voles (*Microtus oeconomus*), reduce photoperiodic induced pars tuberalis thyroid-stimulating hormone β-subunit (*Tshβ*) expression to inhibit gonadal development when food is scarce. Reduction in gonadal size is more pronounced in tundra voles, in which the hypothalamic Kisspeptin (*Kiss1*) system seems involved in downregulating gonadal development, especially in males. Low temperature additionally leads to decreased hypothalamic RF-amide related peptide (*Rfrp3*) levels, which may facilitate further suppression of gonadal growth. Shutting off the photoperiodic-axis when food is scarce in spring may be an adaptive response to save energy, leading to delayed reproductive organ development until food resources are sufficient for reproduction, lactation and offspring survival. Defining the mechanisms through which metabolic cues modify photoperiodic responses will be important for a better understanding of how environmental cues impact reproduction.

**Summary statement:** This study provides a better understanding of the molecular mechanism through which metabolic cues can modify photoperiodic responses, to adaptively adjust timing of reproductive organ development

## Introduction

Seasonal mammals time their reproduction such that offspring will be born during the most optimal time of year, when temperatures are rising and food is abundant. Because of the absence of inter-annual variation in photoperiodic cycles, many vertebrates use photoperiod as a reliable cue to synchronize intrinsic annual timing mechanisms controlling seasonal adaptation of physiology and behavior (for review, see Baker, 1938; Nakane and Yoshimura, 2019). In mammals, photoperiodic signals are perceived by retinal photoreception, and converted in the brain into melatonin signals regulating gonadal responses by the so called ‘photoperiodic neuroendocrine system’ (PNES) (for review, see: Dardente et al., 2018; Hut, 2011; Nakane and Yoshimura, 2019).

Animals that experience food scarcity reduce their overall food consumption while foraging activity is increased (van der Vinne et al., 2019). This induces a negative energy balance, in which there is less energy available for reproductive investment, because most energy ingested is needed for body tissue maintenance. Energy balance and reproduction are closely related (Ruffino et al., 2014; Schneider, 2004), but its regulatory mechanisms remain to be disclosed. Seasonally breeding animals, such as voles, may use a combination of photic and non-photic seasonal cues to control reproduction. Environmental factors such as ambient temperature, food availability and its behavioral foraging activity response can all affect energy balance and are expected to be involved in adaptive modification of the photoperiodic response to inhibit or accelerate reproductive development (Caro et al., 2013; Hut et al., 2014).

The neuroanatomical networks that underly integration of energy balance information into the photoperiodic response system is largely unknown. Neurons expressing gonadotropin-releasing hormone (GnRH) are known to be the main drivers of the reproductive axis controlling the release of hormones (i.e. LH, FSH) from the pituitary gland (Guillemin, 1977; Schally et al., 1970). Prior studies suggest that mediobasal hypothalamic (MBH) thyroid hormone triiodothyronine (T_3_), which is increased under long photoperiods, may not act on GnRH neurons directly, but rather via other hypothalamic areas, such as the preoptic area (POA), the dorso-/ventromedial hypothalamus (DMH/VMH) and the arcuate nucleus (ARC), which are involved in metabolic regulation (for review, see Hut et al., 2014). Neurons located in those hypothalamic regions communicate directly with GnRH neurons (Hileman et al., 2011), and express RF-amides: Kisspeptin (*Kiss1*) (Smith et al., 2005a; Smith et al., 2005b) and RF-amide related peptide (*Rfrp3*), which are candidates for integrating environmental cues, mediating seasonal reproductive responses (Janati et al., 2013; Klosen et al., 2013; Revel et al., 2008). Kiss1 functions as a strong activator of GnRH neurons, and therefore is an important regulator of puberty onset and reproduction (De Roux et al., 2003; Seminara et al., 2004).

To investigate mechanisms of adaptive modification of the photoperiodic response, we decided to use two vole species as study organisms (common vole, *Microtus* arvalis, Pallas 1778 and tundra vole, *Microtus oeconomus*, Pallas 1776). Voles may be the ideal species to study these questions since voles can have strong photoperiodic responses and a functional canonical PNES system (Król et al., 2012; van Rosmalen et al., 2020). On the other hand, voles are also known to be able to respond to the environment in a more opportunistic way (Daketse and Martinet, 1977; Ergon et al., 2001; Negus and Berger, 1977; Nelson et al., 1983; Sanders et al., 1981). common voles are distributed in central Europe (38-62°N), whereas tundra voles are distributed at more northern latitudes (48-72°N). Voles from our two lab populations originate from the same latitude in the Netherlands (53°N), which is for the common vole at the center of its latitudinal range, and for the tundra vole at the southern boundary of its latitudinal range. For this reason, it is expected that our common vole lab population is better adapted to the local environment at 53°N than our tundra vole lab population, which may be better adapted to more northern latitudes. At northern latitudes, tundra voles live under isolating snow covers for a substantial part of the year, which may make photoperiod an unreliable cue for seasonal adaptation in this species. Presumably differences in hypothalamic neurobiological mechanisms may underly the different breeding strategies of the common and the tundra vole.

In this study, voles were exposed to photoperiodic transitions mimicking spring, under which both ambient temperature and food availability was manipulated. By implementing the work-for-food (WFF) paradigm we can induce different levels of natural food scarcity in the laboratory leading to a negative energy balance in small rodents on a high workload (Hut et al., 2011; van der Vinne et al., 2014). We assessed how (which genes), and where in the brain photoperiodic and metabolic cues are integrated to mediate reproductive responses, and how this neurobiological system is differently shaped in two closely related vole species.

## Materials and Methods

### Animals

All experimental procedures were carried out according to the guidelines of the animal welfare body (IvD) of the University of Groningen, and all experiments were approved by the Centrale Comissie Dierproeven of the Netherlands (CCD, license number: AVD1050020186147). common voles, *Microtus arvalis* (Pallas 1778) were obtained from the Lauwersmeer area (Netherlands, 53° 24’ N, 6° 16’ E, (Gerkema et al., 1993). Tundra or root voles, *Microtus oeconomus* (Pallas 1776) were obtained from four different regions in the Netherlands (described in van de Zande *et al*., 2000). All voles in this study were indoor-bred as an outbred colony at the University of Groningen.

All voles used in this study were gestated and born under a short photoperiod (SP, 8h light:16h dark) and transferred to a long photoperiod at the day of weaning at either 21±1°C or 10±1°C. Based on photoperiodic dose-response-curves for gonadal mass from our prior research (van Rosmalen *et al*., in preparation), we selected the photoperiod were maximum gonadal responses were reached, to obtain identical physiological status for both species at the start of the experiments (common voles: 16h light: 8h dark; tundra voles: 14h light:10h dark). Animals were transferred to cages (12.5 × 12.5 × 20 cm) provided with running wheels (14 cm diameter) when they were 35 days old. *Ad libitum* food was available for all animals until they were 40 days old. Animals were provided with water *ad libitum* throughout the course of the experiments.

### Work-for-food protocol

In the work-for-food protocol (starting when animals were 40 days old), animals had to make a set number of wheel revolutions in order to receive a 45 mg grain based food pellet (630 J per pellet) (F0165; Bio-Serv, Flemington NJ, USA), using a computer controlled food dispenser (Med Associates Inc., St.Albans VT, USA). All animals started on a low workload protocol (100 revolutions/ pellet = 0.03 m/J), which is similar to *ad libitum* food conditions, since there were always pellets present in the cages. Half of the animals were subsequently exposed to an increasing workload paradigm in which workload was increased daily by an additional 10-30 revolutions per pellet. Detailed description of the work-for-food protocol for this experiment was published elsewhere (van Rosmalen and Hut, in review). In short, the increase in workload per day was titrated to obtain moderate individual body mass loss (0 - 0.5 gram/day) and the amount of earned pellets per day (>44 kJ/day). All voles were weighed every other day throughout the course of the experiments, in order to carefully monitor growth and to keep animals above 75% of their initial body mass (35 days old).

### Tissue collections

Animals were sacrificed by decapitation, with prior CO2 sedation, 17 ±1 hours after lights OFF at 75 days old. During brain dissection, special care was taken to include the intact pituitary stalk containing the pars tuberalis by cutting the stalk half way between hypothalamus and pituitary gland residing in the hypophysial fossa. To further dissect the hypothalamus, the cerebellum, bulbus olfactorius and frontal cortex were removed by coronal cuts. A sagittal hypothalamic tissue block was obtained by lateral sagittal cuts at the hypothalamic sulci and a horizontal cut at 4 mm distance from the ventral border of the hypothalamus. The posterior hypothalamus (containing pars tuberalis, DMH, VMH and Arcuate nucleus) and anterior hypothalamus (containing POA, PVN, PVz and SCN) were separated by a coronal cut at posterior border of the optic chiasm and the mammillary bodies. After dissection, it was checked that the remainder of the pituitary gland, containing the pars nervosa, pars intermedialis and pars distalis, was still intact in the sella turcica covered by dura mater. The extracted tissues were flash frozen within 2-4 minutes after death in liquid N2 and stored at - 80°C until RNA extraction. Reproductive organs were dissected and cleaned of fat, and wet masses of testes, paired ovary and uterus were measured (±0.0001 g).

### RNA extractions/ Reverse Transcription and Real-time quantitative PCR

Total RNA was extracted from the anterior and posterior hypothalamic area using TRIzol according to the manufacturer’s protocol (Invitrogen, Carlsbad, CA, USA). Frozen pieces of tissue (~20 mg) were homogenized in 0.5 ml TRIzol in a TissueLyser II (Qiagen, Hilden, Germany) (2 × 2 min at 30 Hz) using tubes containing a 5 mm RNase free stainless-steel bead. 0.1 ml chloroform was added for phase separation. Following RNA precipitation by 0.25 ml of 100% isopropanol, the obtained RNA pellets were washed with 0.5 ml of 75% ETOH. RNA was diluted in RNase-free H2O (range 20-100 μl) and quantified on a Nanodrop 2000 spectrophotometer (Thermo Scientific, Waltham, MA, USA). Subsequently, DNA was removed by a DNase I treatment (Invitrogen, Carlsbad, CA, USA), and equal RNA quantities were used for cDNA synthesis by using RevertAid H minus first strand cDNA synthesis reagents (Thermoscientific, Waltham, MA, USA). Reverse transcription (RT; 20 μl) reactions were prepared using 1 μg RNA, 100 μmol 1-1 Oligo(dT)18, 5x Reaction buffer, 20 U μl-1 RiboLock RNase Inhibitor, 10 mmol 1^−1^ dNTP Mix, RevertAid H Minus M-MuLV Reverse Transcriptase (200 U μl^−1^) (Table S1). RNA was reversed transcribed by using a thermal cycler (S1000; Bio-Rad, Hercules, CA, USA). Incubation conditions used for RT were: 45°C for 60 minutes followed by 70°C for 5 minutes. Transcript levels were quantified by real-time qPCR using SYBR Green (KAPA SYBR FAST qPCR Master Mix, Kapa Biosystems). 20 μl reactions were carried out in duplo for each sample by using 96-well plates in a Fast Real-Time PCR System (StepOnePlus; Applied Biosystems, Waltham, MA, USA) (Table S2). Primers for genes of interest were designed using Primer-BLAST (NCBI). All primers were designed based on the annotated *Microtus ochrogaster* genome (NCBI:txid79684, GCA_000317375.1), and subsequently corrected for gene specificity in the genomes of the common vole, *Microtus arvalis* (NCBI:txid47230, GCA_007455615.1), and the tundra vole, *Microtus oeconomus* (NCBI:txid64717, GCA_007455595.1), which both have been sequenced as a collaborative effort between the Hut-lab (Groningen), the Hazlerigg-lab (Tromsø), and the Sandve-lab (Oslo) (Table S3). Relative mRNA expression levels were calculated based on the ΔΔCT method using *Gapdh* as reference gene (Pfaffl 2001).

### Statistical analysis

Sample size (N=6-8) was determined by a power calculation (α=0.05, power=0.95) based on the effect size (d=2.26) of our previous study, in which gonadal mass was assessed in female and male voles under two different photoperiods (van Rosmalen et al., 2020). Effects of workload, temperature and interactions on gonadal weight, body mass and gene expression levels were determined using type I two-way ANOVAs. Tukey HSD post-hoc pairwise comparisons were used to compare groups. Statistical significance was determined at *α =* 0.05. Statistical results can be found in the supplementary information (Table S4). Analyses were performed using RStudio (version 1.2.1335) (R Core Team, 2013), and all figures were generated using the ggplot2 package (Wickham, 2016).

## Results

### Food scarcity reduces reproductive organ mass even under long photoperiods

The reduced body mass in high workload voles (Fig. 1G-J, Table S4) confirms that a negative energy balance was induced by the ‘work-for-food’ protocol. This negative energy balance also caused a 15-47% reduction in testes mass (Fig. 1A,D; Table S4), a 0-50% reduction in ovary mass (Fig. 1B,E; Table S4) and a 18-60% reduction in uterus mass (Fig. 1C,F; Table S4). This effect appeared to be stronger in tundra voles, and in female voles at low temperature.

**Figure 1.**
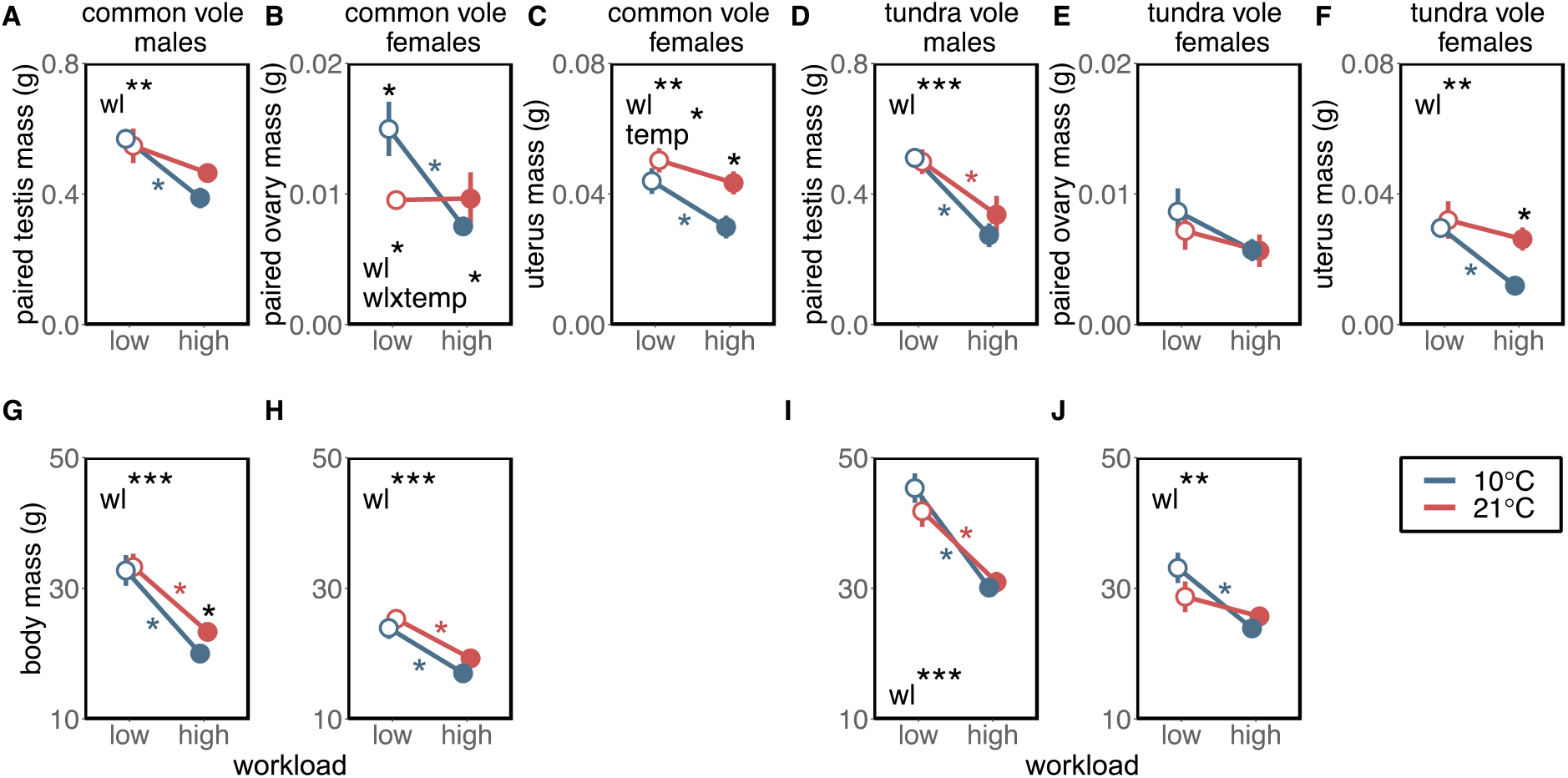
Food scarcity and ambient temperature effects on gonadal weight and body mass in male and female voles. (A,D) paired testis mass, (B,E) paired ovary mass, (C,F) uterus mass and (G-J) body mass for common and tundra voles respectively at low (open symbols) or high workload (filled symbols), at 10°C (blue) or 21°C (red). Data are presented as means ± s.e.m. (n = 6-8). Significant effects (two-way ANOVA) of workload (wl), temperature (temp) and interactions (wlxtemp) are shown: *p < 0.05, **p < 0.01, ***p < 0.001. Significant differences between groups (one-way ANOVA) are indicated by asterisks. Statistic results for ANOVAs can be found in Table S4.

### Food scarcity under long photoperiod suppresses Tshβ expression in the pars tuberalis

To test at what level of the signaling cascade metabolic cues act to modify PNES output signals, we measured gene expression levels in the posterior and anterior hypothalamus. Because the posterior hypothalamic block did not contain the pars distalis, *Tshβ* expression can be exclusively attributed to the pars tuberalis. *Tshβ* expression was significantly reduced (50% reduction) in male voles at high workloads (Fig. 2A,C; Table S4). In males, this effect is stronger in common voles than in tundra voles (Fig. 2A,C, Table S4). In females, this effect was only observed in common voles at 10°C (Fig. 2B,D; Table S4). Temperature did not affect *Tshβ* expression, but a significant reduction was found only in female common voles at 10°C (Fig. 2A-D; Table S4). Overall, *Tshβ* and gonadal weight show a positive relationship (Fig. 3A-D), indicating that *Tshβ* is involved in suppressing gonadal development when food is scarce.

**Figure 2.**
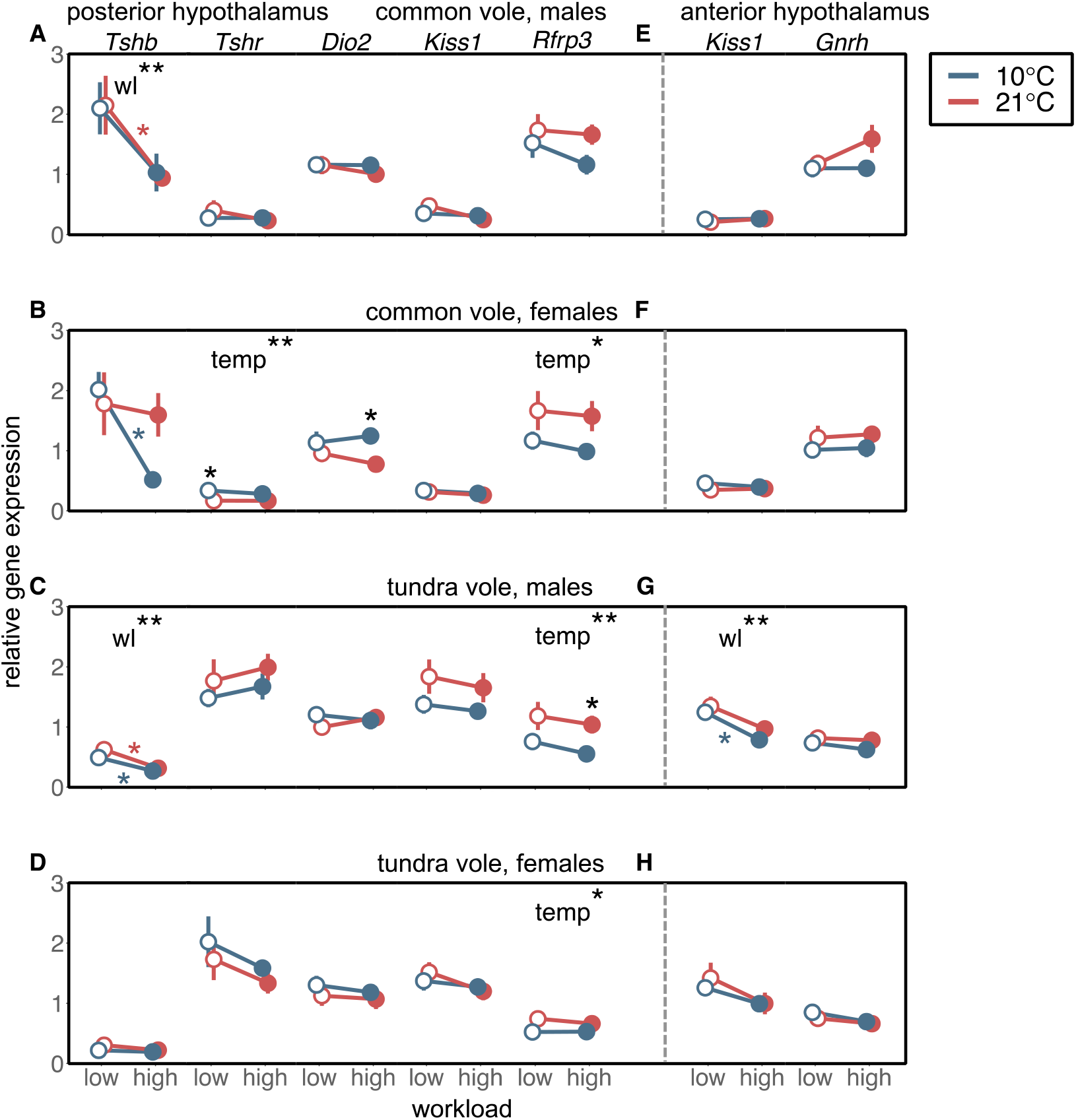
Food scarcity and ambient temperature affect gene expression in the posterior and anterior hypothalamus. Relative gene expression levels of Tshβ, Tshr, Dio2, Kiss1 and Rfrp3 in the posterior hypothalamus and Kiss1 and Gnrh in the anterior hypothalamus for (A, E) common vole males, (B, F) common vole females, (C, G) tundra vole males and (D, H) tundra vole females respectively, at low (open symbols) or high workload (filled symbols), at 10°C (blue) or 21°C (red). Data are presented as means ± s.e.m. (n = 6-8). Significant effects (two-way ANOVA) of workload (wl) and temperature (temp) are shown: *p < 0.05, **p <0.01, ***p < 0.001. Significant differences between groups (one-way ANOVA) are indicated by asterisks. Statistic results for ANOVAs can be found in Table S4.

**Figure 3.**
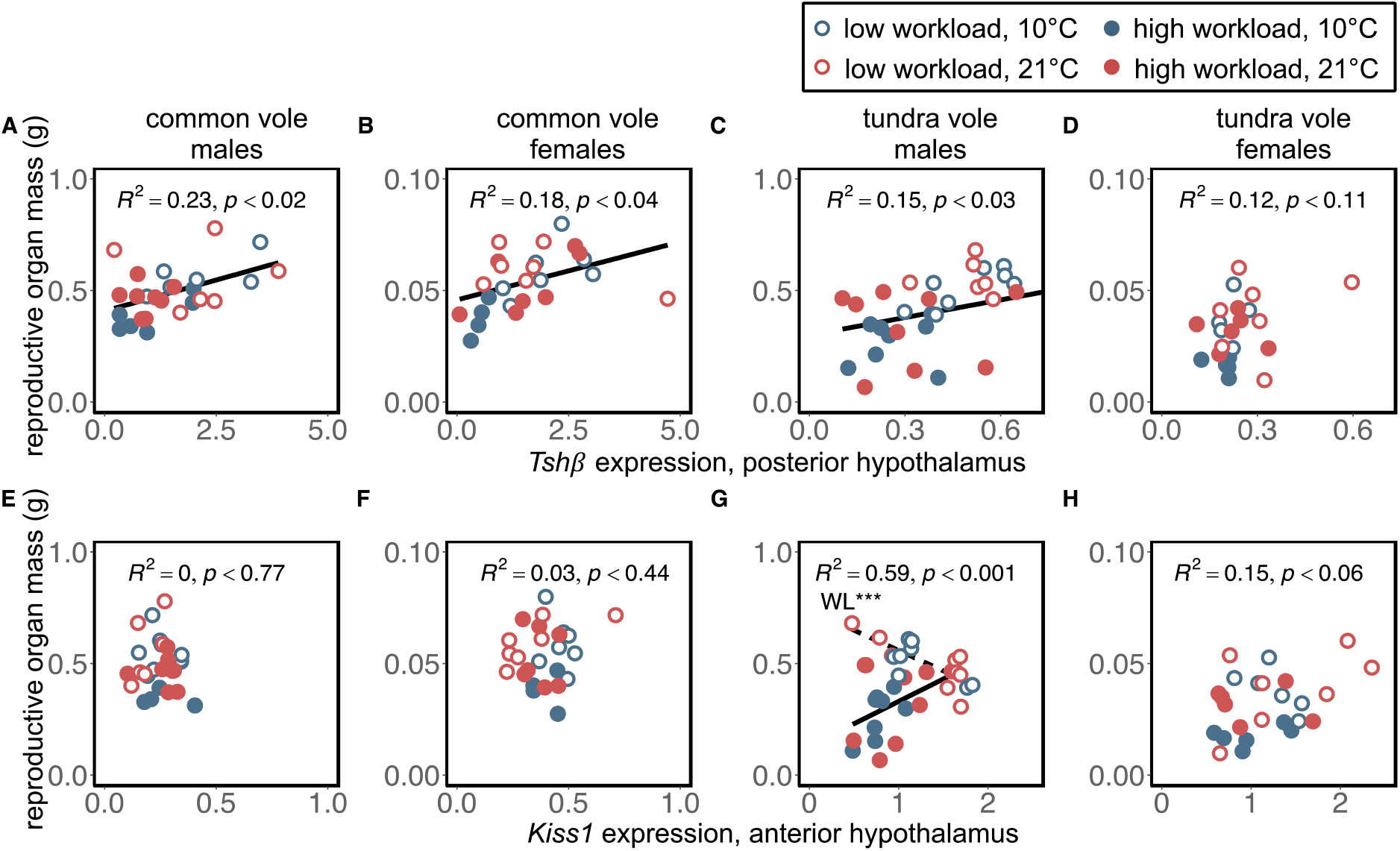
Relationship between reproductive organ mass, Tshβ and Kiss1 expression. Correlations between (A-D) Tshβ expression in the posterior hypothalamus and reproductive organ mass (male: paired testis weight, females: paired ovary + uterus weight), and between (E-H) Kiss1 expression in the anterior hypothalamus and reproductive organ mass for common vole males, common vole females, tundra vole males and tundra vole females respectively at low (open symbols) or high workload (filled symbols), at 10°C (blue) or 21°C (red). Linear models are fitted and R2 and p-values are shown.

After translation, TSHβ and αGSU locally dimerize to form active TSH, which can bind to its receptor (TSHr) located in the tanycytes around the third ventricle of the brain. Workload did not affect *Tshr* expression in both sexes of both species (Fig. 2A-D, Table S4). Although common vole females show slightly elevated *Tshr* expression at 10°C, general *Tshr* levels were lower in common voles than in tundra voles (Fig. 2A-D, Table S4).

Although TSH generally leads to increased *Dio2* levels, workload induced changes in *Tshβ* are not reflected in *Dio2* expression (Fig. 2A-D), suggesting that modifying factors other than TSH can affect posterior hypothalamic *Dio2* expression.

Kiss1 expressing neurons are located in the ARC nucleus in the posterior hypothalamus and are involved in metabolic regulation and food intake. For this reason, it was unexpected that workload and temperature did not affect *Kiss1* expression in the posterior hypothalamus (Fig. 2A-D, Table S4).

The DMH and VMH of the posterior hypothalamus are nuclei that are involved in the regulation of feeding behavior and both these areas are capable of expressing *Rfrp3*. Although overall *Rfrp3* expression was higher at 21°C, no effects of workload on *Rfrp3* expression were observed (Fig. 2A-D, Table S4).

### Food scarcity suppresses Kiss1 signaling in the anterior hypothalamus of tundra vole males

Kiss1 expressing neurons in the anterior hypothalamus are located in the POA, which is involved in temperature regulation (Hrvatin et al., 2020; Takahashi et al., 2020). *Kiss1* expression in the anterior hypothalamus is decreased at high workload in tundra vole males (Fig. 2G, Table S4), whereas *Kiss1* is close to zero in all groups of common voles (Fig. 2E, F; Table S4). Surprisingly, *Kiss1* expression in the anterior hypothalamus was not affected by temperature (Fig. 2E-H, Table S4). Anterior hypothalamic *Kiss1* shows a positive relationship with gonadal weight only in tundra vole males at high workload (Fig. 3E-H). This finding indicates that the *Kiss1* system is involved in modifying photoperiodic responses when food is scarce in tundra voles, but not in common voles.

Perikarya of GnRH neurons are also located in the POA and increased frequency of pulsatile release of GnRH through axonal projections into the median eminence is described to regulate gonadotropin release in the pars distalis of the pituitary gland (Lincoln and Fraser, 1979). Surprisingly, *Gnrh* expression in the anterior hypothalamus was not affected by workload nor temperature (Fig. 2E-H, Table S4).

Fitted linear models revealed that reproductive organ mass can be best predicted by: *Tshβ* and *Tshr* in common vole males (*F*2,23 = 7.44, *p* < 0.01); *Tshβ* in common vole females (*F*1,23 = 4.89, *p* < 0.05); *Tshβ, Rfrp,* anterior hypothalamic *Kiss1* in tundra vole males (*F*3,28 = 8.47, *p* < 0.001); *Tshβ* and anterior hypothalamic *Kiss1* in tundra vole females (*F*2,21 = 3.84, *p* < 0.05).

## Discussion

Our data demonstrate that photoperiodic responses driving gonadal activation can be modified by negative energy balance. Food scarcity seems to act in part via the pars tuberalis to downregulate local levels of TSH, which leads in turn to suppression of gonadal growth, especially in common voles. tundra vole males additionally seem to use the hypothalamic *Kiss1* system to control reproduction when food is scarce at long photoperiods. Kiss1 is one of the main drivers for GnRH neuron activation (De Roux et al., 2003; Han et al., 2005; Han et al., 2015; Seminara et al., 2004). It is therefore surprising that observed patterns in *Kiss1* expression are not reflected in GnRH expression. Although, reproductive organ mass in tundra vole females is reduced at high workloads, no effects at the level of candidate genes have been observed. In general, low temperature enhances the inhibitory effects of reduced energy intake on gonadal size. Within the hypothalamus we show that reduced *Rfrp3* levels may cause decreased reproductive organ mass observed at low temperature.

Here, we chose to investigate the reproductive effects of a negative energy balance in young animals, since voles can reach sexual maturity within 40 days depending on environmental conditions. A negative energy balance under long photoperiod exerts similar effects on testis size of common (Fig. 1A) and tundra voles (Fig. 1D) as in Deer mice, *Peromyscus maniculatus* (Nelson et al., 1997). The lack of this effect in Siberian hamsters, *Phodopus sungorus*, may be explained by the fact that food was minimally reduced to 80-90% of *ad libitum* in this study (Paul et al., 2009). Photoperiod seems to be the driving factor for gonadal development in animals under positive or neutral energy balance, or even with a moderate negative energy balance. A further reduction in food intake counteracts the stimulating effects of long photoperiods on gonadal development, leading to small testes, ovaries and uterus (Fig. 1). Although, large testes are generally associated with high spermatogenic activity and high androgen levels, vole testicular weight may drop in summer while spermatogenic activity remains high (Adams et al., 1980; Wang et al., 2019). Therefore, testicular weight is a reliable indicator for fertility in spring, but less so in summer. Here, voles were exposed to spring photoperiod transitions, therefore, we assume that in our study testes mass is a reliable predictor for fertility. Temperature did not affect testicular weight under long photoperiods when food is available *ad libitum* (Fig. 1A,D). This finding is consistent with prior reports in Siberian hamsters (Steinlechner et al., 1991) and Prairie voles, *Microtus ochrogaster* (Nelson et al., 1989).

Lowering ambient temperature under high feeding related workloads further increases metabolic demands in females, as confirmed by reduced uterine size (Fig. 1C,F). Ovaries of Syrian hamsters (*Mesocricetus auratus*) at low temperature or in the absence of light did not change in size, but had fewer follicles and corpora lutea (Reiter, 1968). This indicates that ovary mass is a bad indicator for hormonal secretory activity. Here, we did not perform histological analysis on ovaries, therefore, this data should be interpreted with caution. On the other hand, small uteri at low temperature are related to reduced height of secretory epithelium and the number of endometrial glands (Reiter, 1968). This confirms that the decline in uterine weight at high workload and low temperature, is the result of incomplete development of uterine glands. This may lead to infertility, since uterine glands are essential for pregnancy (Cooke et al., 2013).

Energetically challenged voles do not enter torpor, as observed in house mice (Hut et al., 2011), but average body temperature is decreased by ~0.5°C, yielding limited energy savings (Nieminen *et al.* 2013; van der Vinne *et al.* 2015; van Rosmalen and Hut, in review). This results in reduced reproductive investment, because all ingested energy is needed for maintaining organ function crucial to survive. Reproductive organ development may persist as ambient temperature and food resources are sufficient for lactation and pup growth.

Our data show that hypothalamic gene expression is related to modify the PNES axis under energetically challenging conditions to reduce gonadal activation. The short duration of pineal melatonin release under long photoperiods lead to increased pars tuberalis *Tshβ*, which serves a pivotal role in the PNES (Hanon et al., 2008; Ono et al., 2008). The present study reveals that the photoperiodic induced *Tshβ* signal can be downregulated by a negative energy balance in common vole males at both temperatures, common vole females only at 10°C and in tundra vole males at both temperatures (Fig. 2A,B,C). Thus, reduced food availability decreases *Tshβ* mRNA at the level of the pars tuberalis, either by decreasing transcription or by increasing post-transcriptional processes. This indicates that a negative energy balance can modify photoperiodic responses at the level of the pars tuberalis or even more upstream in the photoperiodic-axis, primarily in common voles.

Pars tuberalis derived TSH binds locally to TSH receptors (TSHr) in the tanycytes, where it systematically leads to increased DIO2 (Guerra et al., 2010; Hanon et al., 2008; Nakao et al., 2008). The observed *Tshβ* suppression caused by a negative energy balance is not reflected in tanycyte *Dio2* expression (Fig. 2A-D). On one hand, this suggests that TSH modulates central T3 levels and ultimately gonadal development, via pathways parallel to the DIO2/DIO3 system. On the other hand, sex steroid feedback on gene expression in the tanycytes, but not in the pars tuberalis, as observed in ewes, could provide an explanation for unaltered *Dio2* levels (Lomet et al., 2020). The two-fold higher *Dio2* levels in this study compared to our previous experiments might be explained by the fact that here animals were born at short photoperiod and transferred to long photoperiod at weaning, whereas our previous study used constant long photoperiod conditions (van Rosmalen et al., 2020). This effect of maternal photoperiodic programming on tanycyte gene expression has previously been confirmed (Sáenz de Miera et al., 2017; van Rosmalen et al., in preparation). In addition, two to three-day fasted rats show elevated *Dio2* mRNA levels in tanycytes (Coppola et al., 2005; Diano et al., 1998). This might be an acute effect, which disappears when food is restricted for longer periods as in our study (i.e. 35 days). Stable *Tshβ* and *Dio2* levels at different temperatures at low workload under long photoperiods in spring programmed voles are confirmed by in situ hybridization in our prior experiments (van Rosmalen et al., in preparation). This indicates that our brain dissections in combination with RT-qPCR are a reliable method to assess gene expression at the level of the pars tuberalis and the tanycytes.

At the level of the posterior hypothalamus, where the DMH/VMH are located, low temperature induced a small reduction in *Rfrp3* (i.e. *Npvf*) expression, but consistent with Siberian hamsters (Paul et al., 2009), no effect of food scarcity was detected (Fig. 2A-D). In seasonal rodents, *Rfrp3* synthesis appeared to be primarily regulated by photoperiod (for review, see Angelopoulou et al., 2019), but here we show that *Rfrp3* may also be an important regulator of the PNES to adaptively respond to ambient temperature changes. This finding is consistent with previous reports, showing that *Rfrp3* is a hypothalamic biomarker of ambient temperature, independent of nutritional status in mice (Jaroslawska et al., 2015). Moreover, our findings are consistent with a field study in wild Brandt’s voles, *Lasiopodomys brandtii*, in which elevated *Rfrp3* levels were observed during the warmest part of the year (June-August) (Wang et al., 2019). Since RFRP3 is expected to mediate reproductive-axis function, our findings suggest that downregulation of *Rfrp3* by low temperature may be responsible for decreased reproductive organ mass observed at low temperature under high feeding related workloads (Fig. 1A-F).

We next focused on the hypothalamic Kiss1-system, because of its potential role in the integration of photoperiodic and metabolic cues controlling reproductive activity (Caro et al., 2013; Hut et al., 2014; Simonneaux, 2020). Hypothalamic Kisspeptin neurons act on GnRH neurons driving gonadotropin release which promotes gonadal development (for review, see Simonneaux, 2020). *Kiss1* expression in the posterior hypothalamus, where the ARC is located, was not affected by either food or temperature (Fig. 2A-D). ARC *kiss1* expression can be reversed by strong negative sex steroid feedback (Greives et al., 2008; Rasri-Klosen et al., 2017; Sáenz De Miera et al., 2014), which may explain similar *Kiss1* and *Gnrh* levels in different experimental groups (Fig. 2). In Siberian hamsters, food restriction causes a decrease in ARC *Kiss1* expression (Paul et al., 2009). Since whole coronal sections were used in the study of Paul et al. 2009, thalamic and cortical areas contribute to the detected *Kiss1* signal, whereas we exclusively used hypothalamic tissue.

It seems important to note that the role of Kisspeptin in PNES regulation may be less generalizable over different species. For instance, reversed photoperiodic effects on ARC *Kiss1* expression have been shown in Syrian versus Siberian hamsters (Klosen et al., 2013). Furthermore, hypothalamic *Kiss1* expression in common voles is extremely low (Fig. 2A, B, E, F). These findings suggest that Kisspeptin systems have species-specific functions in regulating reproduction. Although *Kiss1* in the posterior hypothalamus was not affected, food scarcity caused downregulation of *Kiss1* expression in the anterior hypothalamus of tundra vole males (Fig. 2G). Although, this effect may be caused by positive sex steroid feedback on POA *Kiss1* expression (Ansel et al., 2010), direct effects of metabolic cues cannot be excluded. Contrary to our expectations, anterior hypothalamic *Kiss1* was not affected by temperature. common voles exhibit extremely low *Kiss1* levels under all conditions. This indicates that the Kiss1-system is not involved in metabolic neuroendocrine control of reproduction in common voles and it is conceivable that *Rfrp3* in this species has taken over the role of *Kiss1*. Interestingly, tundra voles have a functional Kiss-system, which may explain the greater reduction in reproductive organ mass at a negative energy balance. It would be important to investigate how gene-silencing and overexpression (*Tshβ*, *Kiss1* and *Rfrp3*) in specific hypothalamic regions may affect reproductive development, to disentangle causal relationships between hypothalamic gene expression and reproductive responses related to energy balance.

Differences in responses to food scarcity between common and tundra voles, suggest that tundra voles use food as a more important external cue to time reproduction. This seems to be in line with the hypothesis that tundra voles may use an opportunistic breeding strategy, while common voles use breeding strategy that is more driven by photoperiod (van Rosmalen et al., 2020). At northern latitudes, where tundra voles are abundant, voles live for a large part of the year under snow covers where light input is blocked. Reproducing whenever food is available, either as leaves in summer and autumn or as roots during winter and spring, may be a favorable breeding strategy in such environments.

Our findings show that a negative energy balance induced by food scarcity and ambient temperature, modifies long day responses in the PNES: In general, food scarcity regulates the photoperiodic regulated *Tshβ* response in the pars tuberalis (primarily in common voles), and the *Kiss1* response in the anterior hypothalamus (exclusively in tundra vole males), whereas temperature regulates *Rfrp3* in the posterior hypothalamus. Shutting off the photoperiodic-axis when food is scarce or temperatures are low, is an adaptive response that favors individual somatic maintenance and survival at the expense of reproductive organ development and current reproductive output. In addition, delaying reproductive onset will yield energy savings, which results in less required foraging time and reduced exposure to predation, which will further increase individual survival and the probability of successful future reproductive attempts (van der Vinne et al., 2019). Defining the mechanisms through which metabolic cues modify photoperiodic responses will be important for a better understanding of how annual cycling environmental cues shape reproductive function and plasticity in life history strategies.

## List of abbreviations

ARC: arcuate nucleus
*Dio2*: iodothyronine deiodines 2
DMH/VMH: dorso-/ventromedial hypothalamus
Gnrh: gonadotropin-releasing hormone
*Kiss1*: kisspeptin
PNES: photoperiodic neuroendocrine system
POA: preoptic area
*Rfrp3* (*Npvf*): RF-amide related peptide
*Tshβ*: thyroid-stimulating hormone β-subunit
*Tshr*: thyroid-stimulating hormone receptor

## Acknowledgements

We thank L. de Wit for help during tissue collections, G.J.F. Overkamp for technical assistance, Prof.dr. M.P. Gerkema for establishing the common vole colony and Dr. C. Dijkstra and Dr. L. van de Zande for establishing the tundra vole colony at the University of Groningen.

## Author contributions

LvR and RAH designed the research, LvR performed the research and analyzed the data, LvR and RAH wrote the manuscript.

## Data availability

The data that support the findings of this study are openly available in Figshare at https://doi.org/10.6084/m9.figshare.13522700.vl.

